# Progeny counter mechanism in malaria parasites is linked to extracellular resources

**DOI:** 10.1101/2023.01.20.524890

**Authors:** Vanessa S. Stürmer, Sophie Stopper, Patrick Binder, Anja Klemmer, Nils B. Becker, Julien Guizetti

## Abstract

Malaria is caused by the rapid proliferation of *Plasmodium* parasites in patients and disease severity correlates with the number of infected red blood cells in circulation. Parasite multiplication within red blood cells is called schizogony and occurs through an atypical multinucleated cell division mode. The mechanisms regulating the number of daughter cells produced by a single progenitor are poorly understood. We investigated underlying regulatory principles by quantifying nuclear multiplication dynamics in *Plasmodium falciparum* and *knowlesi* using super-resolution time-lapse microscopy. This revealed that the number of daughter cells was statistically independent of the duration of the nuclear division phase, which confirms that a counter mechanism, rather than a timer, regulates multiplication. *P. falciparum* cell volume at the start of nuclear division correlated with the final number of daughter cells. As schizogony progressed, the nucleocytoplasmic volume ratio, which has been found to be constant in all eukaryotes characterized so far, increased significantly, possibly to accommodate the exponentially multiplying nuclei. Depleting nutrients by dilution of culture medium caused parasites to produce less merozoites and reduced proliferation but did not affect cell volume or total nuclear volume at the end of schizogony. Our findings suggest that the counter mechanism implicated in malaria parasite proliferation integrates extracellular resource status to modify progeny number during blood stage infection.

## Introduction

Malaria-causing parasites undergo a complex lifecycle with two critical population bottlenecks during transmission from mosquito to human and back. To overcome those bottlenecks the parasite proliferates extensively in the mosquito midgut, the human liver and red blood cells. In these different stages the parasite generates vast numbers of daughter cells within one cell cycle that can range over four orders of magnitude (Matthews et al., 2018). Malaria pathogenesis, which still kills more than 600,000 people per year (WHO, 2021), occurs during the proliferation of *Plasmodium* spp. in the human blood. After red blood cell (RBC) invasion the parasite generates a variable number of daughter cells, called merozoites, that can range from 12 to 30 (Garg et al., 2015; Mancio-Silva et al., 2013; Reilly et al., 2007; Simon et al., 2021b). Understanding regulatory mechanisms of parasite multiplication is important as the number of merozoites influences growth rate and therefore can contribute to virulence (Mancio-Silva et al., 2013). What limits the number of merozoites is largely unclear.

The division mode of *Plasmodium* spp. is called schizogony and diverges significantly from the classical binary fission observed in most model organisms (Fig. 1A). Schizogony entails a particular cell cycle regulation, which proceeds by multiple asynchronous nuclear divisions not interrupted by cytokinesis and leads to a multinucleated schizont stage (Arnot et al., 2011; Francia and Striepen, 2014; Gerald et al., 2011; Gubbels et al., 2021, 2020; Guttery et al., 2022; White and Suvorova, 2018). Only thereafter nuclei are packaged into merozoites during the segmenter stage, which is followed by egress from the host cell (Absalon et al., 2016; Rudlaff et al., 2020). The released merozoites invade a new RBC and start the next intraerythrocytic development cycle (IDC). IDC durations vary between *Plasmodium* species but usually constitute multiples of 24 hours. *P. falciparum*, which causes the most prevalent and severe form of malaria, requires 48 h and produces 20 merozoites on average (Simon et al., 2021b). The IDC of cultured parasites can deviate from this baseline by a few hours (Reilly Ayala et al., 2010; Reilly et al., 2007). After an initial growth phase, DNA replication, which precedes the first nuclear division, starts at about 30 hours post invasion (hpi) in *P. falciparum* (Ganter et al., 2017; McDonald and Merrick, 2022). *P. knowlesi*, a zoonotic malaria parasite mostly found in macaques, is a rising threat to malaria eradication and has a shorter IDC of 24 h (Lee et al., 2022). *P. knowlesi* was recently adapted for in vitro culture, where it produces about 12 merozoites on average (Mohring et al., 2019; Moon et al., 2013; Simon et al., 2021b). Few functional studies have quantified effects on merozoite number (Dorin-Semblat et al., 2013, 2008; Kumar et al., 2014; Mancio-Silva et al., 2013; Morahan et al., 2020; Robbins et al., 2017). The fact that the parasite can adapt its merozoite number is evidenced by the seminal discovery of a nutrient sensing pathway requiring the KIN kinase, which decreases merozoite number upon calorie restriction in mice (Mancio-Silva et al., 2017). What mechanisms define the number of merozoites and the duration of schizogony remains unclear.

**Figure 1.**
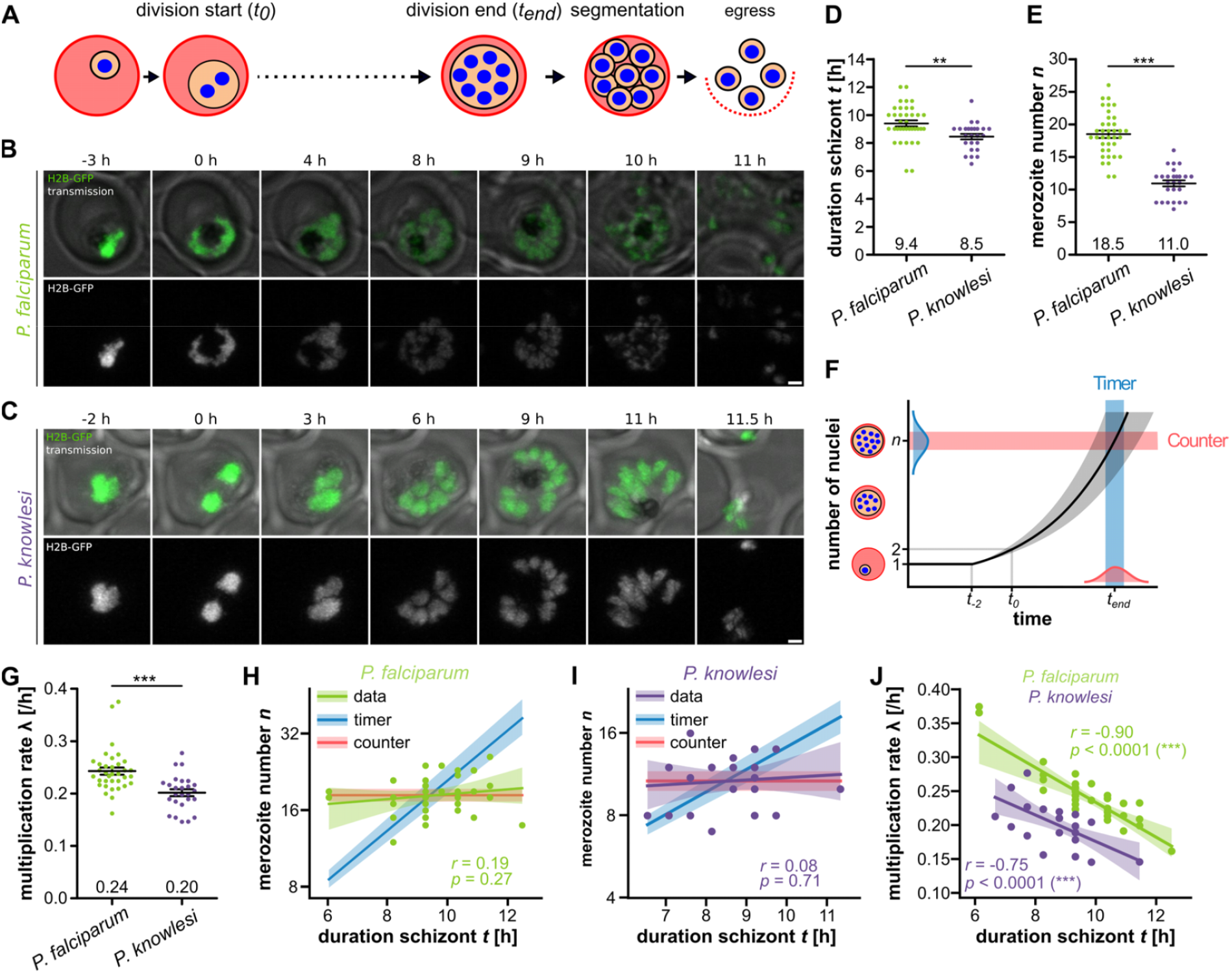
Duration of schizont stage does not correlate with merozoite number in P. falciparum and P. knowlesi. **A** Schematic of schizogony during asexual blood stage development **B** Airyscan-processed time-lapse images of P. falciparum strain 3D7 ectopically expressing H2B-GFP. 0 h indicates first nuclear division. Shown are maximum intensity projections. Scale bars are 1 μm. **C** as in B for P. knowlesi strain A1-H.1. **D** Quantification of duration of schizont stage in hours and **E** merozoite number. Error bars show mean and SEM. All statistical analysis: t-test with Welch’s correction. **F** Schematic of increase of nuclear number throughout schizogony until maximal value (n) is reached at a specific value (counter, red) or after a specific time (blue). **G** Quantification of multiplication rates based on t and n. **H** Plot comparing predicted regression curves for timer (blue) and counter (red) model with data bootstrapped with 95% confidence interval. **I** as in H for P. knowlesi. **J** Bootstrapped regression curves of t against λ. Given are Pearson correlation coefficient r and p values. N = 35 for P. falciparum and N = 26 for P. knowlesi all from three independent replicates.

In eukaryotes, two main phenomenological mechanisms linking cell division to cell growth have been described (Facchetti et al., 2017; Jones et al., 2019). In a timer mechanism, a cell completes division after a preset time has elapsed. While the interdivision time may be stochastic, it is uncorrelated with cell size or growth speed. By contrast, in a sizer mechanism a cell completes division once a specific extensive cellular parameter, such as cell size, has reached a set value. Again, the preset size value may be stochastic but is required to be uncorrelated with interdivision time. In various uni- and multicellular eukaryotes both mechanisms have been described and appear to be implemented by sensing different cellular parameters before engaging in cell division (Cadart et al., 2018; Facchetti et al., 2017; Gao et al., 1997; Li et al., 2016; Neumann and Nurse, 2007; Ondracka et al., 2018).

In the present context of schizogony, the relevant system size is the final number of nuclei, which reflects the number of merozoites produced per invasion event. We previously speculated that either a timer mechanism or a counter mechanism, which is analogous to a sizer, could control merozoite number (Simon et al., 2021b). A timer predicts that duration of schizogony correlates with final number of nuclei, while a counter mechanism would conclude schizogony after a pre-set number of nuclei is formed independently of the required time. Differentiating between those models is critical to guide further analysis as a counter mechanism would imply a molecular factor limiting growth, while a timer mechanism would necessitate a biochemical reaction functioning as a molecular hourglass in a way that is independent of nuclear growth. Analysis of *P. falciparum* DNA replication and nuclear division dynamics observed that the duration of the first nuclear cycle covaries with overall duration of nuclear multiplication. This excluded a timer and thereby confirmed a counter as the mechanism regulating termination of nuclear multiplication (Klaus et al., 2022). A direct correlation between duration of schizont stage and merozoite number is, however, still missing. It is also unclear which cellular or extracellular parameters are relevant for determining final merozoite number.

Most other studies that have quantified schizogony have so far relied on fixed cell or population analysis. Here, we employ time-resolved data of key cellular parameters in individual cells. By using Airyscan-detector-based super resolution live cell imaging modalities we achieve sufficiently low photo-toxicity and high spatial resolution to reliably track live cells throughout schizogony and count individual merozoites. We correlate duration of the schizont stage, cell size, and merozoite number in single cells, which provides evidence for a counter mechanism. We further uncover that throughout schizogony *P. falciparum* infringes on the otherwise ubiquitously constant N/C-ratio (Cantwell and Nurse, 2019), and find that diluting nutrients from culture media reduces progeny number.

## Results

### Nuclear multiplication dynamics confirm counter mechanism in *P. falciparum* and *knowlesi*

To correlate duration of the schizont stage with merozoite number, we generated *P. falciparum* and *P. knowlesi* strains ectopically expressing GFP-tagged Histone 2B (H2B-GFP), which labels nuclear chromatin (Fig. 1B-C). To resolve individual daughter nuclei in live cells, we employed Airyscan detector-based microscopy, which improves spatial resolution by a factor of about 1.7 beyond the diffraction-limit, and recorded schizogony from the first nuclear division until egress from the host cell (Mov. S1-2). Imaging under hypoxia conditions and in riboflavin-free media improved cell viability and reduced photo-toxicity (Zigler et al., 1985). Although separating individual nuclei after the first two rounds of division was challenging due to their spatial proximity, the improvements in resolution and 3D image analysis allowed us to count the final number of nuclei routinely and reliably at the transition into the segmenter stage. The final number of segmenter nuclei always perfectly matched the number of merozoites observed just before egress. Since it was also shown that every nucleus is packaged into an individual merozoite (Absalon et al., 2018, 2016), we will from now on refer to this number as merozoite number (n). In this study, we define the start of the schizont stage (t_0_) by the first appearance of two distinct nuclei and its end (t_end_) as the time when the maximal number of nuclei was first reached (Fig. 1A). The duration of the schizont stage (t) for *P. falciparum* was around 9.4 h (Fig. 1D). Remarkably *P. knowlesi*, which has a much shorter IDC (McDonald and Merrick, 2022; Moon et al., 2013), showed an only slightly reduced duration of the schizont stage of around 8.5 h. The merozoite number, however, was much lower, with n=18.5 and n=11.0, in *P. falciparum* and *knowlesi*, respectively (Fig. 1E), which matched with the previously measured values in fixed cells (Simon et al., 2021b). To further validate our assay, we used an alternative marker, the microtubule live cell dye SPY555-Tubulin, in *P. falciparum* (Mov. S3). Defining the first mitotic spindle extension as t_0_ and appearance of subpellicular microtubules as t_end_, resulted in very similar and reproducible values (Fig. S1). Whether merozoite number is set by a counter or by a timer mechanism can be revealed by the correlation of merozoite number with duration of the schizont stage (Fig. 1F). When a timer concludes schizogony earlier or later, the merozoite number will be reduced or increased, respectively. A counter independently sets merozoite number, so no correlation should be observed in that case. The dynamics of nuclear multiplication follow an exponential growth function (Klaus et al., 2022). Merozoite number (*n*) is therefore determined by the nuclear multiplication rate (λ) and duration of the schizont stage (*t*) by *n*(*t*) = 2 e^Jt^. The multiplication rate (λ) was lower in *P. knowlesi* than in *P. falciparum* (Fig. 1G). To test the counter and timer models, we log-plotted *n* against *t* for *P. falciparum* and *P. knowlesi* (Fig. 1H-I). A timer mechanism would predict that a regression line would approximate the slope given by λ, while for a counter the slope would approximate zero. The slope of the regression curve did not significantly deviate from zero (Fig. 1H-I, S1D). Bootstrapping analysis, as developed previously (Klaus et al., 2022), again confirmed that the counter prediction of near-zero slope falls well within the range supported by the data, while the timer prediction is not compatible. Plotting λ against t showed a significant negative correlation (Fig. 1J). This indicates that parasites employ a higher multiplication rate to reach higher merozoite numbers, keeping the duration of schizogony roughly constant, once more incompatible with a timer mechanism. When we shuffled the data by reassigning λ – *t* pairs randomly and then computed resulting merozoite numbers according to a timer model with exponential growth, this strongly increased their variability (Fig. S2). Thus, the negative correlation between λ and *t* (Fig. 1J) reflects a precisely controlled merozoite number during schizogony. Taken together these data support that the counter mechanism acts through modulation of nuclear multiplication rate.

### Duration of merozoite segmentation is constant between *P. falciparum* and *knowlesi*

To interrogate the duration of merozoite formation we quantified the time from end of schizont stage (t_end_) until egress from the host cell, i.e. the segmentation phase (Fig. 1A). We found it to be identical in both species at around 2.7 h (Fig. 2A). The presence of more nuclei did not prolong the time required for segmentation (Fig. 2B), and neither was segmentation time correlated with the duration of the preceding schizont stage (Fig. 2C). This indicates that the time used for merozoite formation is largely fixed and independent of previous events.

**Figure 2.**
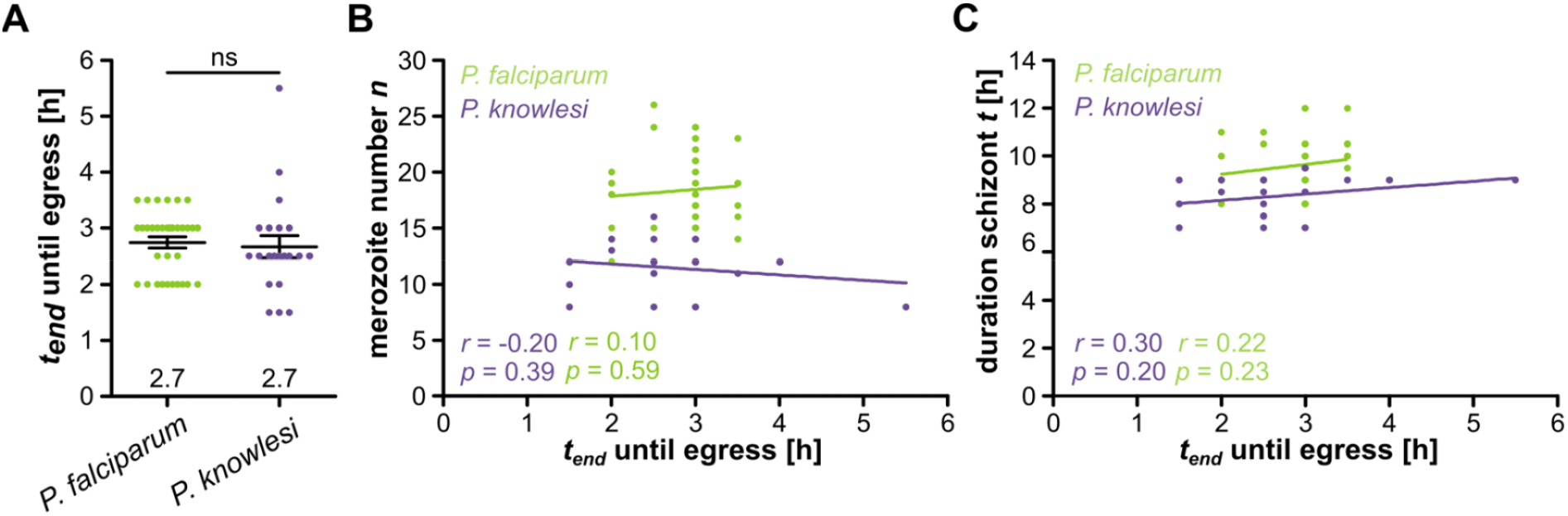
Time required for segmentation does not depend on the number of nuclei. Quantification of time between schizogony end and egress for H2B-GFP expressing P. falciparum (green) and P. knowlesi (blue) in hours. Given are mean and SEM. Statistical analysis: t-test with Welch’s correction. **B** Correlation of merozoite number against segmentation time. **C** Correlation of duration of schizont stage against segmentation time. Given are Pearson correlation coefficient r and p values. N = 33 for P. falciparum and N = 21 for P. knowlesi all from three independent replicates.

### Merozoite number does not depend on host cell diameter

The counter mechanism raises the question which cellular or extracellular parameters might limit merozoite number. Cell size has been identified as an important parameter linked to the initiation of cell division in several organisms (Heald and Gibeaux, 2018; Jones et al., 2019). We therefore investigated whether size of the host erythrocyte might influence merozoite number, possibly by providing more nutrients or more space for the parasite. Since RBC volume is challenging to assess directly in our assay, we measured the average RBC diameter in cells lying flat (Fig. 3A). Host erythrocytes become much more spherical as a consequence of parasite growth (Waldecker et al., 2017). Due to this rounding effect *P. falciparum* and *P. knowlesi*-infected RBCs had a smaller diameter compared to uninfected ones (Fig. 3B). We found no significant correlation between RBC diameter and merozoite number at defined timepoints (Fig. 3C-D). Whether this finding extrapolates to a lack of correlation with RBC volume remains to be investigated.

**Figure 3.**
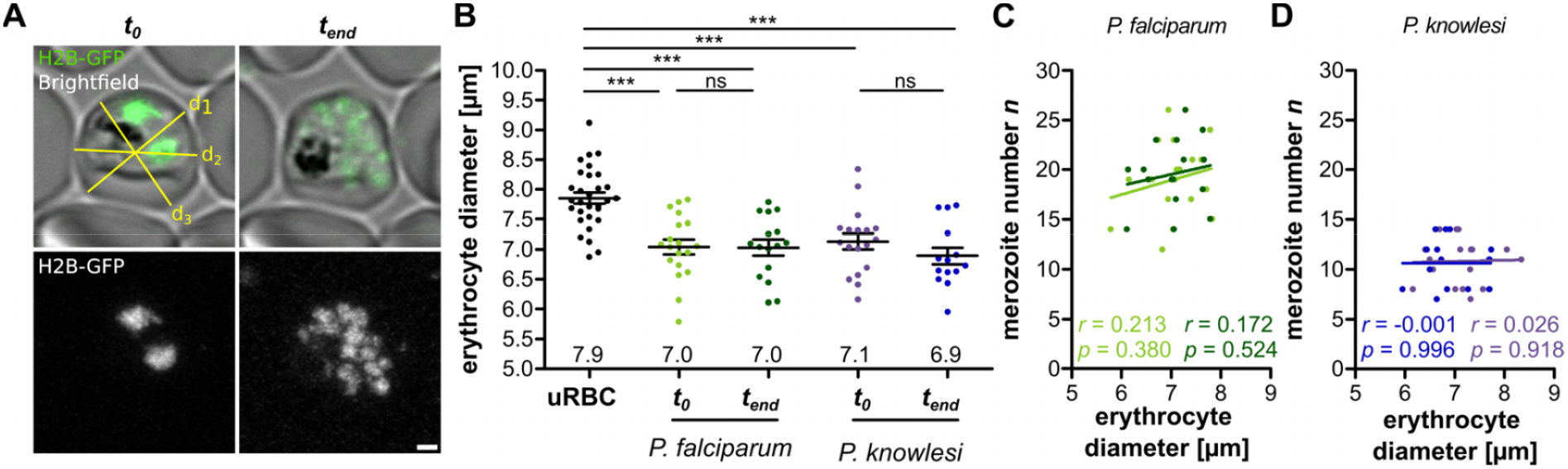
Erythrocyte diameter has no effect on merozoite number in P. falciparum. **A** Airyscan-processed images of P. falciparum 3D7 ectopically expressing H2B-GFP. Erythrocyte diameter was measured by averaging three diameters (yellow lines) at different angles. **B** Erythrocyte diameters in uninfected red blood cells (uRBC) and P. falciparum (green) and P. knowlesi (blue) infected red blood cells. Means of three diameter measurements for each RBC are plotted. Error bars show mean and SEM. Statistical analysis: t-test with Welch’s correction. **C** Correlation of P. falciparum merozoite number against erythrocyte diameter. **D** As in C for P. knowlesi. Given are Pearson correlation coefficient r and p values. N = 30 for uRBC, N = 19 (t_0_) and N = 16 (t_end_) for P. falciparum H2B-GFP and N = 18 (t_0_) and N = 15 (t_end_) for P. knowlesi H2B-GFP all from three independent replicates.

### Cell size at onset of schizont stage correlates with merozoite number

To measure cellular and nuclear volume of the parasite we generated a *P. falciparum* strain ectopically expressing cytosolic GFP and mCherry tagged with a nuclear localization signal (NLS) (Fig. 4A). We could not produce dual-labeled *P. knowlesi* strains due to the lack of an effective second resistance cassette and therefore continued our analysis focusing on *P. falciparum*. The dual-marker *P. falciparum* strain spent the same time in the schizont stage as the H2B-GFP strain although its merozoite number was slightly higher, which could be explained by the absence of histone tagging that might affect nuclear multiplication (Fig. S3A-B). Again, the data clearly supported the counter model (Fig. S3C-D). We acquired long-term time lapse microscopy data of progression through schizogony (Mov. S4) and applied automated image thresholding on the cytosolic GFP signal (Fig. S4) to quantify parasite cell volume (Fig. 4B). These data started up to 7 h before schizont onset and showed a steady increase in total cell volume from ~20 μm^3^ to ~80 μm^3^ around the end of schizogony (Fig. 4C). This aligned with previously reported and modelled values of 18 – 32 μm^3^ at 30 to 34 hpi and 60 – 76 μm^3^ at 48 hpi (Lew et al., 2003; Mauritz et al., 2009; Waldecker et al., 2017). Importantly, we wanted to test whether parasite cell volume did correlate with *n*, λ, and *t*. For analysis we identified three relevant timepoints t_−2_, t_0_, and t_end_. The t_−2_ time point corresponds to the onset of DNA replication which occurs on average 2 h before the first nuclear division (Klaus et al., 2022). We found a significant correlation of merozoite number with the cell volume around the onset of schizogony (Fig. 4D-E), while at the end of schizogony the correlation was not quite significant anymore (Fig. 4F). Consistently, we found a correlation of λ with cell size around the onset of schizogony, while duration of schizont stage was not correlated (Fig. S5). Taken together this suggests that the setting of the counter might correlate with parasite cell size.

**Figure 4.**
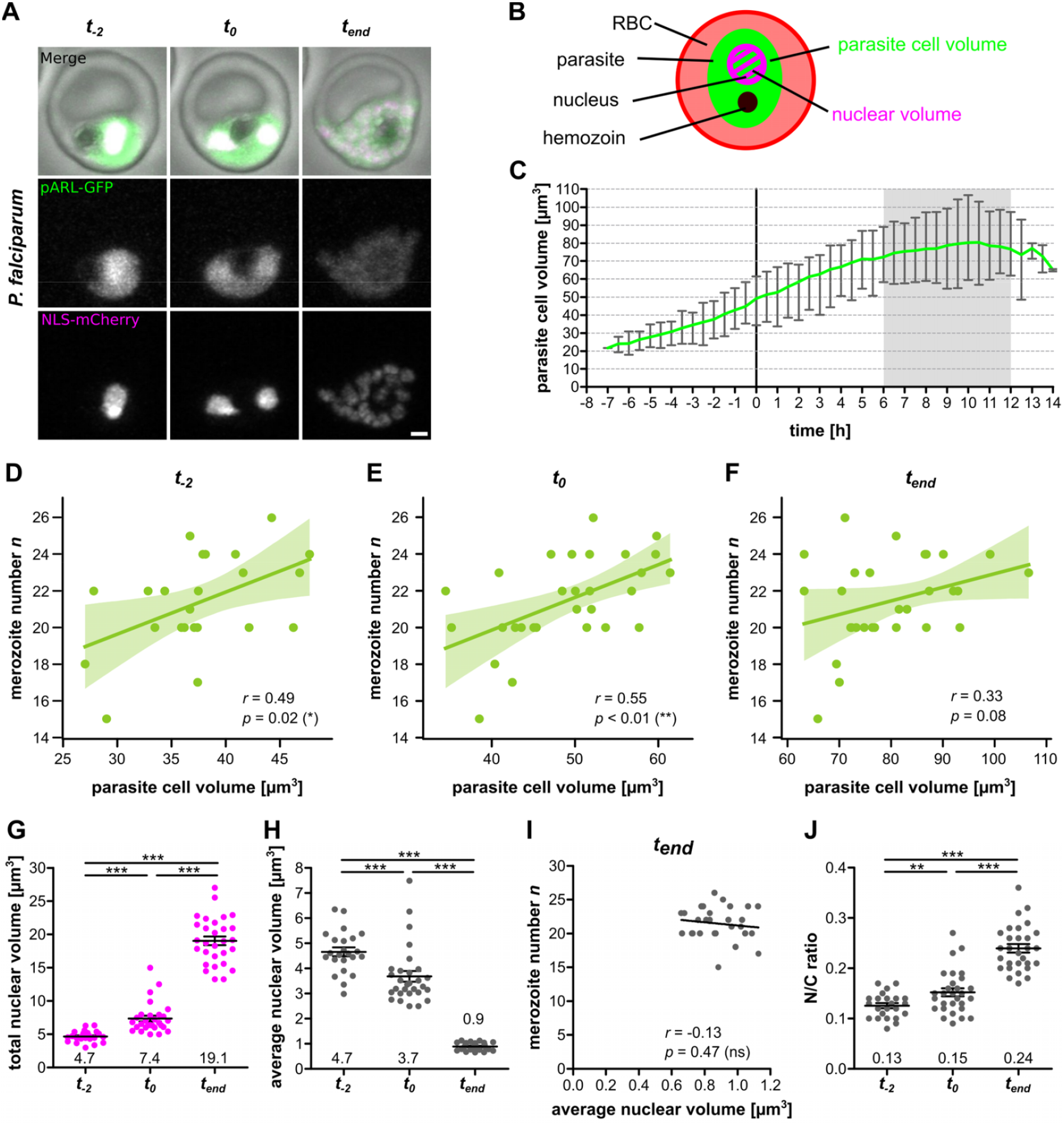
Cell size around schizogony start correlates with merozoite number and N/C-ratio increases. **A** Airyscan-processed time-lapse images of P. falciparum 3D7 ectopically expressing cytosolic GFP (pARL-GFP, green), nuclear mCherry (NLS-mCherry, magenta). Selected timepoints were pre-schizogony (t_−2_), schizogony start (t_0_) and schizogony end (t_end_). Maximum intensity projections are shown. Scale bar is 1 μm. **B** Schematic for segmentation of parasite cell volume (green) and nuclear volume (magenta). **C** Quantification of parasite cell volume over time. Line represents mean and error bars range. Curves were aligned to schizogony start (0 h). Grey zone indicates range of measured schizont stage durations. **D** Correlation of merozoite number against cell volume at pre-schizogony (t_−2_), **E** schizogony start (t_0_) and **F** schizogony end (t_end_). Given are Pearson correlation coefficient r and p values. Values are bootstrapped to 95% confidence interval **G** Total nuclear volume for individual cells at t_−2_, t_0_, and t_end_. **H** Average nuclear volume (total nuclear volume divided by nuclear number) for individual cells. **I** Merozoite number against average nuclear volume at end of schizogony (t_end_). Given are Pearson correlation coefficient r and p values. **J** Total nuclear volume divided by cell volume (N/C ratio) for individual cells. All error bars represent mean and SEM. Statistical analyses: t-test with Welch’s correction. N = 23 for t_−2_ and N = 29 for all others from three independent replicates.

### Nuclear compaction during schizogony does not offset increase of N/C-ratio

Quantification of total nuclear volume was performed by manual thresholding at t_−2_, t_0_, and t_end_, as the decrease of signal over time prevented automated thresholding (Fig. S4). We observed an about 3-fold increase in total nuclear volume throughout the schizont stage (Fig. 4G). Simultaneously the average volume of individual nuclei decreased from 4.7 μm^3^ to 0.9 μm^3^ at the end of schizogony. This indicates that the parasite compacts its nuclei to accommodate their rising numbers within a limited cellular volume. The nuclear volume at t_end_ had a notably low variability (Fig. 4H), and the reduction in nuclear volume did not scale with the total number of nuclei (Fig. 4I), which suggests that the final nuclear volume might represent a state of maximal compaction. Despite nuclear compaction the overall N/C-ratio, which has been described to be universally constant across eukaryotic cell cycles (Cantwell and Nurse, 2019), almost doubled from 0.13 to 0.24 (Fig. 4J), but did not correlate with final nuclear number (Fig. S6). We also approximated nuclear volume in *P. knowlesi* by manual segmentation of the H2B-GFP signal (Fig. 1C), which showed trends similar to *P. falciparum* although the minimal nuclear volume was higher (Fig. S7). Taken together this suggests that the nuclear compaction we document is not a gradual response to the density of nuclei but rather a convergence to a uniformly small nuclear size.

### Nutrient-limited conditions reduce merozoite number

Parasites take up resources for growth from the host red blood cell and the surrounding medium. To test whether nutrient status of the medium affects proliferation we experimented with several dilutions of complete RPMI cell culture medium with physiological 0.9% NaCl solution (Fig. S8). This reduces every media component while maintaining osmolarity and pH. While dilutions to 0.33x or less ultimately caused parasite death, they could be grown over prolonged periods in 0.5x diluted medium (Fig. S8). Hematology analysis showed that RBC indices were nearly identical under all conditions indicating that host cell health is maintained (Fig. S9). Aside a lower parasitemia (Fig. 5A), we could observe a significant reduction in parasite multiplication rate compared to normal 1x culture media condition (Fig. 5B). This was accompanied by a reduced merozoite number as measured by 3D imaging of fixed and Hoechst-stained wild type segmenters (Fig. 5C). Using time-lapse microscopy, we imaged the schizont stage of the cytoplasmic GFP and nuclear mCherry-expressing parasite strain that were switched to 0.5x medium during the preceding ring stage (Mov S5). Schizont stage was slightly longer and the significant reduction in merozoite number was reproducible (Fig. 5D-E). This resulted in a lower multiplication rate (Fig. 5F), while the data remained fully compatible with a counter mechanism (Fig. 5G-H).

**Figure 5.**
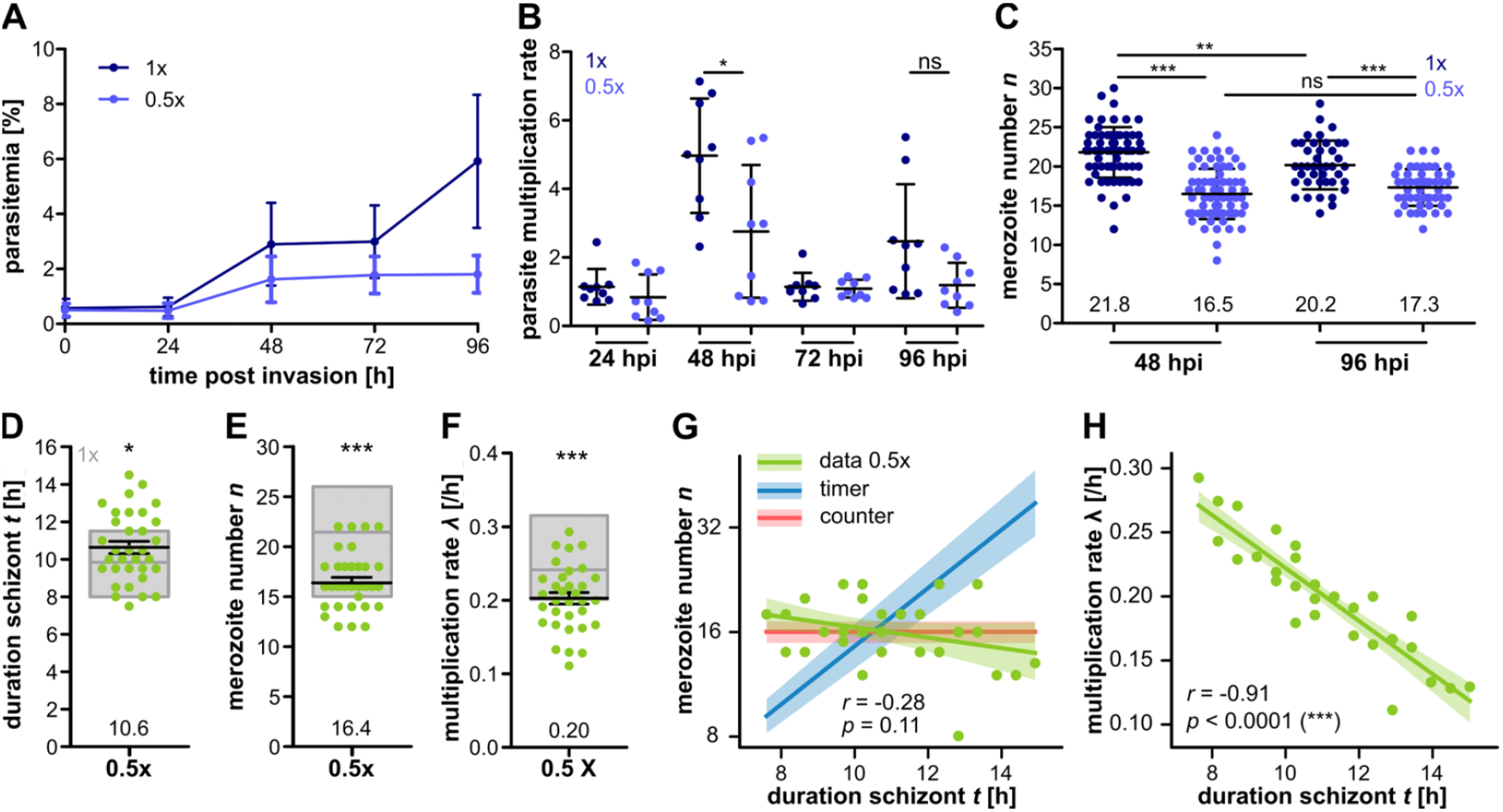
Medium dilution causes reduction in merozoite number in P. falciparum. **A** Growth curve of synchronized 3D7 WT grown in normal 1x (dark blue) and diluted 0.5x (light blue) medium starting as early rings. Medium changed every 24 h. Plotted are mean and SD. **B** Multiplication rate from A. **C** Merozoite numbers counted by fluorescent microscopy after 48 and 96 h. **D** Duration of schizont stage t in hours, **E** merozoite number n and **F** multiplication rate lambda [/h] of P. falciparum 3D7 ectopically expressing cytosolic GFP and nuclear mCherry in 0.5x diluted medium. Parasites were placed in medium conditions 24 - 30 hours prior to imaging. Gray boxes show mean, min and max of previously presented data in 1x conditions (Sup. Fig. 3). **G** Correlation of merozoite number n against schizont duration t and **H** multiplication rate lambda against schizont duration t, data from movies as in D-F. Given are Pearson correlation coefficient r and p values. Values are bootstrapped to 95% confidence interval. Statistical analyses: t-test. N = 3 from three independent replicates for A-B, N= 67 (1x, 48h), 69 (0.5x, 48h), 41 (1x, 96h) and 50 (0.5x, 96h), and N = 33 from ten independent replicates for D – H.

### Nutrient-limited conditions increases average nuclear volume

We also quantified cell volume in 0.5x media conditions and only found a small increase at t_−2_ and t_0_ while at t_end_ cell volumes were similar (Fig. 6A-B). We, however, did not observe a correlation between cell volume and merozoite number at early schizont stages anymore (Fig. 6C-D), while at t_end_ a slight correlation appeared (Fig. 6E). This shows that even though cell size can correlate with merozoite number in standard growth conditions (Fig. 4D-E) the division counter is not strictly coupled to it. A notable difference was the total nuclear volume being higher in 0.5x medium cells at the onset of nuclear division (Fig. 6F). Remarkably, at the end of division it was identic to standard conditions despite the lower final number of nuclei. This translated to a significantly higher average nuclear volume (Fig. 6G), which was, contrary to standard media conditions, anticorrelated with merozoite number (Fig. 6H). Finally, this lower number of nuclei with a higher volume in an equally big cell yielded the same N/C-ratio at t_end_ (Fig. 6I). Despite a resource-limited environment and reduced merozoite number the parasites still converge towards an equivalent total nuclear volume, cell volume, and N/C-ratio (Fig. 6B,F,I). Since these cellular parameters are interlinked, it remains to be seen, which one might be dominant.

**Figure 6.**
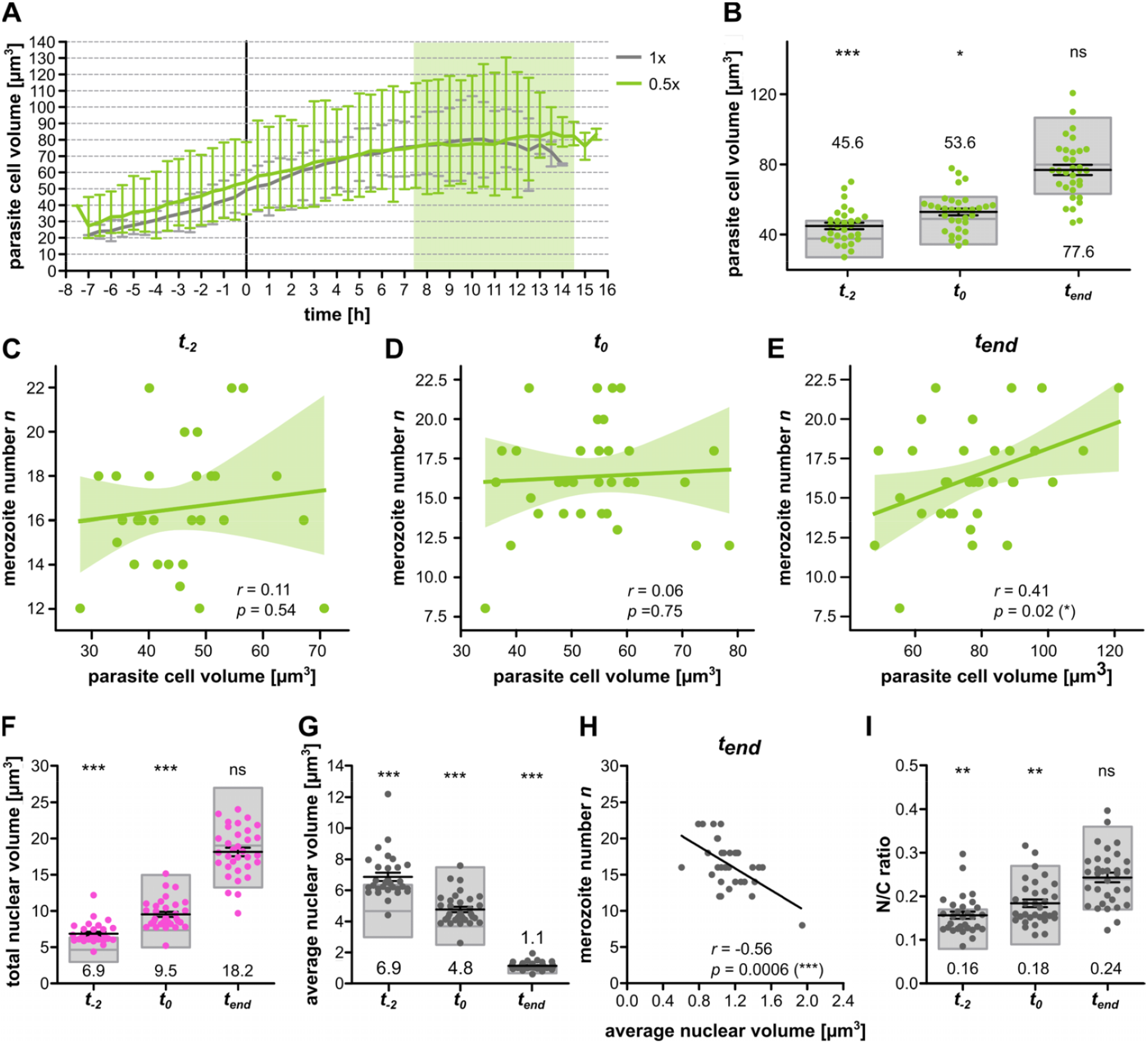
No correlation between cell volume around schizogony start and merozoite number but preserved N/C ratio increase under diluted medium condition. **A** Quantification of parasite cell volume over time of time-lapse imaging with P. falciparum 3D7 ectopically expressing cytosolic GFP and nuclear mCherry in 0.5x diluted medium with 0.9% NaCl. Parasites were placed in diluted medium conditions 24 - 30 hours prior to imaging and late trophozoites were selected for movies. Line represents mean and error bar range; grey line shows data acquired in 1x medium (Fig. 4). Curves were aligned to schizogony start (0 h). Light green zone indicates range of measured schizont stage durations. **B** Comparison of parasite cell volume from A between 0.5x medium (green dots) and 1x medium (grey boxes showing mean, min, max) at pre-schizogony (t_−2_), schizogony start (t_0_) and schizogony end (t_end_). **C** Correlation of merozoite number against cell volume at pre-schizogony (t_−2_), **D** schizogony start (t_0_) and **E** schizogony end (t_end_). Given are Pearson correlation coefficient r and p values. **F** Total nuclear volume for individual cells at t_−2_, t_0_, and t_end_. **G** Average nuclear volume (total nuclear volume divided by nuclear number) for individual cells at timepoints like in F. **H** Merozoite number against average nuclear volume at end of schizogony (t_end_). Given are Pearson correlation coefficient r and p values. **I** Total nuclear volume divided by cell volume (N/C ratio) for individual cells at timepoints described in F. All error bars represent mean and SEM. Statistical analyses: t-test with Welch’s correction. N = 30 for t_−2_ and N = 33 for all others from ten independent replicates.

## Discussion

In this study we quantify key biophysical cell division parameters to correlate duration, cell volume, and nuclear volume with the number of daughter cells generated. Our findings support previous models that parasites use a limiting factor to regulate proliferation (Klaus et al., 2022; Simon et al., 2021b). We also suggest extracellular resources might be linked to modulation of parasite progeny number. While multiplying the nuclei get compacted which reduces the atypical increase in N/C-ratio (Fig. 7). An unexpected finding was that the schizont stage in *P. knowlesi* had a duration similar to *P. falciparum* despite its IDC being much shorter. Recently, the fraction of the schizont stage duration was quantified at about 30% of IDC duration in *P. knowlesi* and *P. falciparum* (McDonald and Merrick, 2022). For *P. knowlesi* the authors found the duration from the first DNA replication, detected by appearance of nuclei pulse-chase labeled with Ethyl-deoxyuridine (EdU), to the peak of median nuclear number in fixed cell populations to be around ~9-11 h. We found 8.5 h on average for the duration of first to last nuclear division, which matches well when adding the 2 h required for the first round of DNA replication (Klaus et al., 2022). For *P. falciparum*, however, the authors found ~15 h, while we estimate the equivalent duration at 11.4 h. Strain to strain variation as well as the different methods used, i.e. time-lapse imaging vs pulse-chase labeling of Thymidine Kinase overexpressing parasites, could explain these differences. Furthermore, the authors indicated a time lag of about 10 h (32 – 42 hpi) between the time point when the first and the last *P. falciparum* parasite starts replication, which limits the temporal resolution of their assay. For their *P. falciparum* strain the authors further indicate an IDC duration of 48 h, while we estimate our strain to be slightly faster.

**Figure 7.**
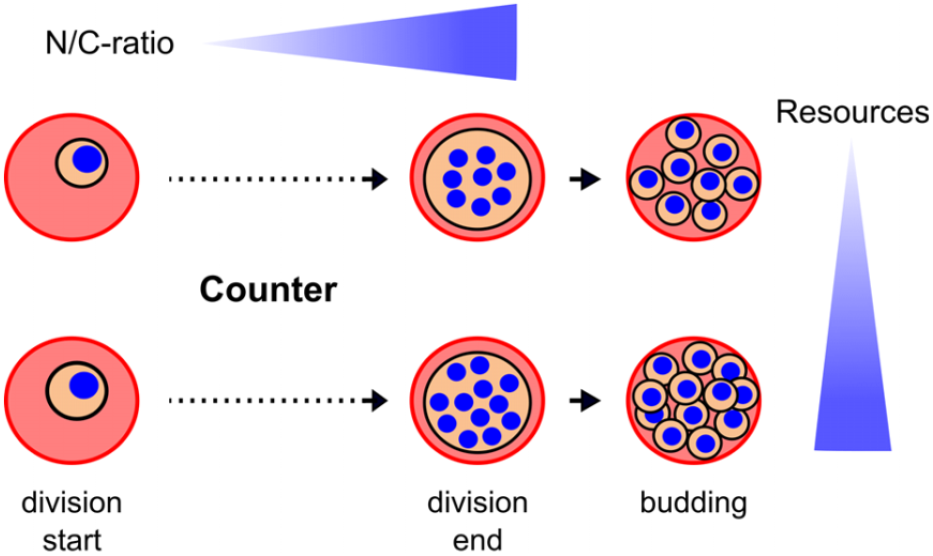
Schematics of nuclear division events leading to high or low numbers of merozoites. As Plasmodium parasites undergo nuclear division, they increase their N/C-ratio. High merozoite numbers can result from larger cells at onset of division but are more clearly linked to nutrient concentration in the cell culture medium.

Why *P. knowlesi* produces less daughter cells in a similar time remains unclear, but one can note that its merozoites are significantly bigger (Lyth et al., 2018), which might limit their number. Among other Plasmodium species, *P. berghei* produces ~12, while *P. chabaudi* only generates ~6-8 merozoites within their 24 h long IDC (Birget et al., 2019). *P. malariae*, whose IDC lasts 72 h, generates only around ~8-10 merozoites per infected RBC, so overall no correlation appears to exist between progeny number and IDC duration across species. For *P. falciparum* and *knowlesi* we, however, consistently observe that the number of merozoites is independent of the time spent on nuclear divisions. These nuclear multiplication data are not compatible with a timer mechanism but support a counter mechanism controlling the completion of schizogony, as has been shown by a previous study analyzing DNA replication dynamics in *P. falciparum* (Klaus et al., 2022). The fact that other multinucleated organisms like the Ichthyosporean *Sphaeroforma arctica* undergo regularly timed nuclear division events that are independent of cell size and growth emphasizes that a counter is not self-evident (Ondracka et al., 2018). Implementing a counter mechanism requires a limiting factor that sets the growth boundary for the system, which we can now attempt to uncover in more detail.

The concept that nutrients are limiting progeny number has already been shown by studying parasite infection in calorie restricted mice (Mancio-Silva et al., 2017). This might provide advantages for the parasite in being able to quickly notice and adapt to changing environmental conditions (Kumar et al., 2021). KIN was identified as regulator mediating between nutrient sensing and transcriptional response. Even though KIN is a divergent kinase it seems to have analogous functions to the TOR or energy sensing pathway, which links nutritional status to cell growth (McLean and Jacobs-Lorena, 2017). This is partly coherent with our finding that cell size around the start of schizogony predicts progeny number. In the green alga *Chlamydomonas reinhardtii,* mother cell size is linked to the number of uniform-sized daughter cells (Li et al., 2016) and is even influenced by light exposure (Heldt et al., 2020). Number of nuclear division events and daughter cell size are dependent on the amount of CDKG1 (D-cyclin-dependent RB kinase) in the mother cell (Li et al., 2016). How cells can sense cell size has been shown in Xenopus where progressive sequestration of importin alpha to the plasma membrane during cell division is used for measuring cell surface-area-to-volume ratios, thereby regulating nuclear scaling (Brownlee and Heald, 2019).

As eukaryotic cells grow their nuclear volume increases to keep the N/C-ratio constant, although it is not clear why. The “Kern-Plasma-relation theory” had already been proposed by Hertwig and Boveri at the beginning of the 20^th^ century (Hertwig, 1903) and has been corroborated in uni- and multicellular eukaryotes ranging from mammals to plants and fungi (Cantwell and Nurse, 2019), which includes multinucleated organisms like *Ashbya gossypii* (Dundon et al., 2016). Hence, our observation that malaria parasites do not maintain their N/C-ratio constant was surprising. A key difference, however, to most other studied organisms is that parasite growth is limited by the host cell of which it usually occupies up to 80% at the end of development (Waldecker et al., 2017). In this context it might be a better strategy to infringe on the N/C-ratio to maximize the number of daughter cells that can be fitted in the host cell. Another strategy could be the further reduction of nuclear volume, which is usually not determined by DNA content (Cantwell and Nurse, 2019). Our observation that Plasmodium nuclei converge to a small but very uniform volume of about 0.9 μm^3^ might however suggest a lower limit for nuclear size imposed by the degree to which DNA can be compacted. Coherently the relative cell size of the parasite does not correlate with the relative DNA content as it does in most other cells, which generates specific challenges with respect to gene dosage (Machado et al., 2021). On the other hand, we saw in *P. knowlesi*, which has a similar sized genome, that the minimal nuclear size was 1.5-fold bigger. An alternative explanation is also suggested by our nutrient depletion experiments where despite producing less nuclei the N/C-ratio is increasing to the same level. In merozoites themselves, whose volume was measured at about 1.7 μm^3^ (Dasgupta et al., 2014), the nuclei occupy an even more significant proportion of the entire cell (Hanssen et al., 2013; Rudlaff et al., 2020). Hence, the increase of the N/C-ratio during schizogony could also be viewed as gradual progression towards generating a highly compacted cell stage specialized for invasion.

Discoveries presented here were made possible by imaging new division markers with super-resolution time-lapse microscopy, while mitigating the effects of light-induced stress. Overall, this study highlights that resolving single-cell dynamics of cellular processes over longer time scales must gain importance in the study of malaria parasite proliferation (De Niz et al., 2016; Gruring et al., 2011; Klaus et al., 2022; Yahiya et al., 2022).

The quantitative and dynamic analyses carried out here provide important insights into the regulation of progeny number and provides a biophysical and cell biological framework for further quantitative analysis of the rapid proliferation of malaria-causing parasites in the blood.

## Materials and Methods

### Parasite cultivation and growth curves

All *P. falciparum* cell lines were cultured in 0+ erythrocytes in cRPMI (RPMI 1640 containing 0.2 mM hypoxanthine, 25 mM HEPES, 0.5% Albumax and 12.5 μg/ml Gentamycin) and maintained at a hematocrit of 2.5% and a parasitemia of 1-5%. Dishes were kept incubated at 37°C with 90% humidity, 5% 0_2_ and 3% CO_2_. *P. knowlesi* cell lines (WT kindly provided by Robert Moon (Mohring et al., 2019; Moon et al., 2013)) were cultured as described above but cRPMI was additionally supplemented with 10% horse serum. For diluted media conditions the cRPMI as described above was diluted with 0.9% NaCl solution to the indicated final medium concentration and afterwards sterile filtered. Parasite growth for all growth curves was determined using Giemsa-stained thin blood smears. At least 1000 Erythrocytes or 20 parasites were counted for calculating the parasitemia. Double infections were counted as single infections. Parasites were provided with fresh media every 24 or 48 h for unsynchronized and synchronized growth curves respectively. Synchronization to a window of 3-4 h was achieved using pre-synchronization with 0.5% sorbitol followed by MACS purification and reinvasion.

### Cloning of pARL-H2B-GFP and pPkconcrt-H2B-GFP

To generate pARL-H2B-GFP, pARL-GFP was digested with XhoI and AvrII for linearization. PfH2B was amplified from gDNA with forward primer TTCTTACATATAActcgagATGGTATCAAAAAAACCAGC and reverse primer GAACCTCCACCTCCcctaggTTTGGAGGTAAATTTTGTTA. Ligation was performed using Gibson Assembly (NEB, E2611S) (Gibson et al., 2009) and the resulting plasmid was checked using Sanger sequencing.

To generate Pkconcrt-GFP, an episomal construct for expression of GFP-tagged proteins, we used Pkcon-GFP (kindly provided by Robert Moon) which was digested with XmaI and NotI-HF to cut out the PkHSP70 promoter. PkCRT promoter was amplified from gDNA with forward primer TATAGAATACTCGCGGCCGCGGTAACGGTTCTTTTTCTGA and reverse primer CCCTTGCTCACCATCCCGGGTGTGTGTTTGTTTTAGCGGTG. Ligation was performed using Gibson Assembly and the resulting plasmid was checked using Sanger sequencing with forward primer CACAGGAAACAGCTATGACC and reverse primer GGTGGTGCAGATGAACTT. To generate the pPkconcrt-H2B-GFP plasmids for episomal expression of PkH2B, pPkconcrt-GFP was linearized using XmaI and PkH2B was amplified from gDNA with forward primer TAAAACAAACACACACCCGGGATGGTATCCAAAAAGCCAGCG and reverse primer CATAGAACCTCCACCTCCCCTAGGCTTGGAGGTAAATTTCGTAACGGC. Ligation was performed using Gibson Assembly. Resulting plasmids were checked by a control digest using XmaI and AvrII and sequenced using Sanger sequencing with forward primer CGATGCGGTAGAAGAGTCGT and reverse primer CTTGTGGCCGTTTACGTCGC.

### *P. knowlesi* transfection

Parasites were synchronized prior to transfection by using 55% Nycodenz cushions and reinvasion assay. 20 μg of plasmid per reaction was precipitated using 1/10 vol. 3M sodium acetate and 2 vol. cold 100% ethanol, washed with cold 70% ethanol and resuspended in sterile TE buffer. For transfection segmenters were enriched using 50 mM cyclic GMP-dependent protein kinase (PKG) inhibitor ML10 and subsequently transfected by electroporation. Selection for plasmid was done using 2.5 nM WR99210 (Jacobus Pharmaceuticals) 24 hours after transfection.

### *P. falciparum* transfection

pARL-H2B-GFP was transfected into 3D7 WT parasites by electroporation of sorbitol synchronized ring-stage parasites and pARL-GFP (kindly provided by Jude Przyborski) into parasites already ectopically expressing NLS-mCherry (previously provided by Markus Ganter (Simon et al., 2021a)) with 50–100 μg of purified plasmid DNA (QIAGEN). Plasmid precipitation was done prior to transfection as stated above. To select for the plasmid, we used 2.5 nM WR99210 (Jacobus Pharmaceuticals) 24 hours after transfection.

### Preparing cells for live cell imaging

For live cell imaging, resuspended parasite culture was washed twice with pre-warmed iRPMI, and seeded on μ-Dish (ibiTreat, 81156) or μ-Dish 35 mm Quad (ibiTreat, 80416) for 10 min at 37°C similar as described before (Gruring et al., 2011; Mehnert et al., 2019). Cells were washed with iRPMI until a monolayer remained. For H2B-GFP movies μ-Dishes were filled completely with imaging medium, meaning phenol red-free RPMI1640 with stable Glutamate and 2 g/L NaHCO_3_ (PAN Biotech, P04-16520) otherwise supplemented like stated above and previously equilibrated in the incubator beforehand for several hours. Dishes were sealed with parafilm and kept at 37°C prior of imaging. For all other movies 600 μl per well of imaging medium lacking Riboflavin (Biomol, R9001-04.10) but otherwise supplemented to the same composition as stated above was added in the 35 mm Quad dish and the lid was placed loose on top, as the imaging chamber was incubated with 5% 0_2_ and 5% CO_2_ during the complete imaging period. In case of *P. knowlesi* movies 10% horse serum was added additionally to the imaging medium. For live-Tubulin staining SPY555-Tubulin or SPY650-Tubulin (Spirochrome) was added to the imaging medium to a final concentration of 500 nM. For diluted media conditions cells were transferred into 0.5x media dilution 24 - 30h before imaging, and live cell imaging was performed in 0.5x imaging medium (imaging medium diluted as described above) selecting for late trophozoites to ensure altered medium conditions since early ring stage for respective parasites.

### Super-resolution live cell imaging

Live cell imaging was performed using point laser scanning confocal microscopy on a Zeiss LSM900 microscope equipped with the Airyscan detector using Plan-Apochromat 63x/1,4 oil immersion objective. The imaging chamber was incubated at 37°C and 5% 0_2_ and 5% CO_2_ in a humidified environment. Images were acquired at multiple positions using an automated stage and the Definite Focus module for focus stabilization. Images were taken at an interval of 30 min – 1 hour for a total period of around 20 hours. Multichannel images were acquired sequentially in the line scanning mode using 488 nm, 561 nm, and 640 nm diode lasers at 0.1 % laser power except for volume movies at 0.3%. Brightfield images were obtained from a transmitted light PMT detector. GaAsP PMT and Airyscan detectors were used with the gain adjusted at 500 - 900V. Image size was typically 20.3 × 20.3 μm with a pixel size of 0.04 μm. Z-stack slices were imaged at an interval of 0.35 nm for a total range of 6 – 7 μm. Subsequently, ZEN Blue 3.1 software was used for the post 3D Airyscan processing with automatically determined default Airyscan Filtering (AF) strength.

### Image analysis and statistics

Image analysis was performed with Fiji. Counting of nuclei was done manually in 3D (z-stack) using the segmenter stage where nuclei were separated the most. Start of schizogony was determined when two separate nuclei were visible for the first time. End of schizogony was determined by no more appearing nuclei and beginning of segmentation. Merozoite number was confirmed by blinding. Movies were acquired in three or more independent replicates for all experiments. Prism5 was used for all statistical analysis. R-values indicate how well x and y correlate. P-values of correlation analysis indicate whether the slope of regression curve is significantly non-zero. T-tests were performed for statistical tests between two conditions and in case of unequal variances with Welch’s correction. To determine the range of regression slopes compatible with data in Fig.1HIJ, we performed bootstrap re-sampling from the data, and report the resulting (2.5, 97.5) percentile range of linear regression slopes. To investigate the importance of λ-t correlations appearing in Fig.1J, we randomly reassigned the (λ,t) data points, generating samples of uncorrelated synthetic data; for each such data point we calculated a merozoite number as predicted by a timer mechanism, shown in Fig. S2.

### Volume determinations

Volume determinations were performed using Fiji. For cellular volume, GFP signal in all z-slices per timepoints was thresholded using the automated thresholding method “RenyiEntropy”. Particles with a smaller area than 0.05 μm^2^ were excluded as well as holes to eliminate the area occupied by hemozoin. The volume was calculated by the slicing method approximating a solid volume by slicing it in regular intervals, adding up areas of all sections and multiplying it with the slicing interval. Therefore, the total GFP area per timepoint was multiplied with the z interval of 0.35 μm for calculating the final volume. Image analysis macro used for processing can be provided upon request. For nuclear volume, mCherry signal was manually thresholded at timepoints t_−2_, t_0_ and t_end_ by setting a visual minimal intensity value. The subsequent volume analysis was done as stated above.

### Automated hematology analysis of erythrocytes

0+ erythrocytes have been kept in different medium dilutions (1x, 0.5x, 0.33x and 0.25x) at a hematocrit of 5% for 24 and 48 h prior to automated determination of red blood cell indices using a Sysmex XP-300. Hematocrit was adjusted to physiological levels before analysis. All conditions have been tested in triplicates with blood from one donor.

### Merozoite counting in fixed segmenters

Tightly synchronized late-stage parasites grown in 1X and 0.5X medium since early rings were collected at the end of the first and second cycle and enriched in segmenter stages using the cyclic GMP-dependent protein kinase (PKG) inhibitor ML10 at a final concentration of 25 nM for 2 – 3 hours. Cells were seeded on imaging dishes and fixed with 4% paraformaldehyde for 20 min at 37°C. Nuclei were stained using Hoechst. Imaging of individual segmenters was performed using point laser scanning confocal microscopy on a Zeiss LSM900 microscope equipped with the Airyscan detector using Plan-Apochromat 63x/1,4 oil immersion objective. Images were acquired in the line scanning mode using 405 nm diode laser at 0.3 % laser power. Brightfield images were obtained from a transmitted light PMT detector. GaAsP PMT and Airyscan detectors were used with the gain adjusted at 500 – 900V. Image size was 10.1 × 10.1 μm with a pixel size of 0.04 μm. Z-stack slices were imaged at an interval of 0.14 nm for a total range of 8 μm. Subsequently, ZEN Blue 3.1 software was used for the post 3D Airyscan processing with automatically determined default Airyscan Filtering (AF) strength. Merozoites were counted by eye using Fiji and confirmed by blinding. Multiple infections were excluded. The experiment was performed in three technical replicates, counting a minimum of 20 segmenters per replicate.

## Author Contributions

Experiments were planned and designed by V.S.S and J.G. V.S.S carried out the experiments and analysis with the help of S.S and under supervision by J.G. A.K. helped produce transgenic *P. knowlesi* lines. N.B.B and P.B carried out detailed statistical analysis. V.S.S generated figures with help by J.G. J.G. and V.S.S wrote the manuscript with input by P.B. and N.B.B.

## Acknowledgments

We thank: The Infectious Diseases Imaging Platform for imaging support (idip-heidelberg.org). Markus Ganter for providing the NLS-mCherry strain and critical comments on the manuscript. Robert Moon for providing *P. knowlesi* acceptor strain with the Pkcon-GFP plasmid.

## Data and Material Availability Statement

All data is provided in the manuscript or supplemental information. Materials can be provided upon request to the corresponding author.

## References

Absalon S, Blomqvist K, Rudlaff RM, DeLano TJ, Pollastri MP, Dvorin JD. 2018. Calcium-dependent protein kinase 5 is required for release of egress-specific organelles in Plasmodium falciparum. MBio 9:e00130–18. doi:10.1128/mBio.00130-18

Absalon S, Robbins JA, Dvorin JD. 2016. An essential malaria protein defines the architecture of blood-stage and transmission-stage parasites. Nat Commun 7:11449. doi:10.1038/ncomms11449

Arnot DE, Ronander E, Bengtsson DC. 2011. The progression of the intra-erythrocytic cell cycle of Plasmodium falciparum and the role of the centriolar plaques in asynchronous mitotic division during schizogony. Int J Parasitol 41:71–80. doi:10.1016/j.ijpara.2010.07.012

Birget PLG, Prior KF, Savill NJ, Steer L, Reece SE. 2019. Plasticity and genetic variation in traits underpinning asexual replication of the rodent malaria parasite, Plasmodium chabaudi. Malar J 1:222. doi:10.1186/s12936-019-2857-0

Brownlee C, Heald R. 2019. Importin α Partitioning to the Plasma Membrane Regulates Intracellular Scaling. Cell 176:805–815.e8. doi:10.1016/j.cell.2018.12.001

Cadart C, Monnier S, Grilli J, Sáez PJ, Srivastava N, Attia R, Terriac E, Baum B, Cosentino-Lagomarsino M, Piel M. 2018. Size control in mammalian cells involves modulation of both growth rate and cell cycle duration. Nat Commun 9:3275. doi:10.1038/s41467-018-05393-0

Cantwell H, Nurse P. 2019. Unravelling nuclear size control. Curr Genet 65:1281–1285. doi:10.1007/s00294-019-00999-3

Dasgupta S, Auth T, Gov NS, Satchwell TJ, Hanssen E, Zuccala ES, Riglar DT, Toye AM, Betz T, Baum J, Gompper G. 2014. Membrane-wrapping contributions to malaria parasite invasion of the human erythrocyte. Biophys J 107:43–54. doi:10.1016/j.bpj.2014.05.024

De Niz M, Burda P-C, Kaiser G, del Portillo HA, Spielmann T, Frischknecht F, Heussler VT. 2016. Progress in imaging methods: insights gained into Plasmodium biology. Nat Rev Microbiol 15:37–54. doi:10.1038/nrmicro.2016.158

Dorin-Semblat D, Carvalho TG, Nivez M-P, Halbert J, Poullet P, Semblat J-P, Goldring D, Chakrabarti D, Mehra P, Dhar S, Paing MM, Goldberg DE, McMillan PJ, Tilley L, Doerig C. 2013. An atypical cyclin-dependent kinase controls Plasmodium falciparum proliferation rate. Kinome 1:4–16. doi:10.2478/kinome-2013-0001

Dorin-Semblat D, Sicard A, Doerig Caroline, Ranford-Cartwright L, Doerig Christian. 2008. Disruption of the PfPK7 gene impairs schizogony and sporogony in the human malaria parasite Plasmodium falciparum. Eukaryot Cell 7:279–285. doi:10.1128/EC.00245-07

Dundon SER, Chang SS, Kumar A, Occhipinti P, Shroff H, Roper M, Gladfelter AS. 2016. Clustered nuclei maintain autonomy and nucleocytoplasmic ratio control in a syncytium. Mol Biol Cell 27:2000–2007. doi:10.1091/mbc.E16-02-0129

Facchetti G, Chang F, Howard M. 2017. Controlling cell size through sizer mechanisms. Curr Opin Syst Biol 5:86–92. doi:10.1016/j.coisb.2017.08.010

Francia ME, Striepen B. 2014. Cell division in apicomplexan parasites. Nat Rev Microbiol 12:125–136. doi:10.1038/nrmicro3184

Ganter M, Goldberg JM, Dvorin JD, Paulo JA, King JG, Tripathi AK, Paul AS, Yang J, Coppens I, Jiang RHY, Elsworth B, Baker DA, Dinglasan RR, Gygi SP, Duraisingh MT. 2017. Plasmodium falciparum CRK4 directs continuous rounds of DNA replication during schizogony. Nat Microbiol 2:17017-. doi:10.1038/nmicrobiol.2017.17

Gao FB, Durand B, Raff M. 1997. Oligodendrocyte precursor cells count time but not cell divisions before differentiation. Curr Biol 7:152–155. doi:10.1016/S0960-9822(06)00060-1

Garg S, Agarwal S, Dabral S, Kumar N, Sehrawat S, Singh S. 2015. Visualization and quantification of Plasmodium falciparum intraerythrocytic merozoites. Syst Synth Biol 9:23–26. doi:10.1007/s11693-015-9167-9

Gerald N, Mahajan B, Kumar S. 2011. Mitosis in the human malaria parasite plasmodium falciparum. Eukaryot Cell 10:474–482. doi:10.1128/EC.00314-10

Gruring C, Heiber A, Kruse F, Ungefehr J, Gilberger TW, Spielmann T. 2011. Development and host cell modifications of Plasmodium falciparum blood stages in four dimensions. Nat Commun 2:165. doi:10.1038/ncomms1169

Gubbels MJ, Coppens I, Zarringhalam K, Duraisingh MT, Engelberg K. 2021. The Modular Circuitry of Apicomplexan Cell Division Plasticity. Front Cell Infect Microbiol 11:670049. doi:10.3389/fcimb.2021.670049

Gubbels MJ, Keroack CD, Dangoudoubiyam S, Worliczek HL, Paul AS, Bauwens C, Elsworth B, Engelberg K, Howe DK, Coppens I, Duraisingh MT. 2020. Fussing About Fission: Defining Variety Among Mainstream and Exotic Apicomplexan Cell Division Modes. Front Cell Infect Microbiol 5:269. doi:10.3389/fcimb.2020.00269

Guttery DS, Zeeshan M, Ferguson DJP, Holder AA, Tewari R. 2022. Division and Transmission: Malaria Parasite Development in the Mosquito. Annu Rev Microbiol 76:113–134. doi:10.1146/annurev-micro-041320-010046

Hanssen E, Dekiwadia C, Riglar DT, Rug M, Lemgruber L, Cowman AF, Cyrklaff M, Kudryashev M, Frischknecht F, Baum J, Ralph SA. 2013. Electron tomography of Plasmodium falciparum merozoites reveals core cellular events that underpin erythrocyte invasion. Cell Microbiol 15:1457–1472. doi:10.1111/cmi.12132

Heald R, Gibeaux R. 2018. Subcellular scaling: does size matter for cell division? Curr Opin Cell Biol 52:88–95. doi:10.1016/j.ceb.2018.02.009

Heldt FS, Tyson JJ, Cross FR, Novák B. 2020. A Single Light-Responsive Sizer Can Control Multiple-Fission Cycles in Chlamydomonas. Curr Biol 30:634–644.e7. doi:10.1016/j.cub.2019.12.026

Hertwig R. 1903. Ueber die Korrelation von Zell-und Kerngrösse und ihre Bedeutung für die Geschlechtliche Differenzierung und die Teilung der Zelle. Biol Cent 23:49–62.

Jones AR, Band LR, Murray JAH. 2019. Double or Nothing? Cell Division and Cell Size Control. Trends Plant Sci 24:1083–1093. doi:10.1016/j.tplants.2019.09.005

Klaus S, Binder P, Kim J, Machado M, Funaya C, Schaaf V, Klaschka D, Kudulyte A, Cyrklaff M, Laketa V, Höfer T, Guizetti J, Becker NB, Frischknecht F, Schwarz US, Ganter M. 2022. Asynchronous nuclear cycles in multinucleated Plasmodium falciparum facilitate rapid proliferation. Sci Adv 8:1–13. doi:10.1126/sciadv.abj5362

Kumar M, Skillman K, Duraisingh MT. 2021. Linking nutrient sensing and gene expression in Plasmodium falciparum blood-stage parasites. Mol Microbiol 115:891–900. doi:10.1111/mmi.14652

Kumar P, Tripathi A, Ranjan R, Halbert J, Gilberger T, Doerig C, Sharma P. 2014. Regulation of Plasmodium falciparum development by calcium-dependent protein kinase 7 (PfCDPK7). J Biol Chem 289:20386–20395. doi:10.1074/jbc.M114.561670

Lee WC, Cheong FW, Amir A, Lai MY, Tan JH, Phang WK, Shahari S, Lau YL. 2022. Plasmodium knowlesi: the game changer for malaria eradication. Malar J. doi:10.1186/s12936-022-04131-8

Lew VL, Tiffert T, Ginsburg H. 2003. Excess hemoglobin digestion and the osmotic stability of Plasmodium falciparum - Infected red blood cells. Blood 101:4189–4194. doi:10.1182/blood-2002-08-2654

Li Y, Liu D, López-Paz C, Olson BJ, Umen JG. 2016. A new class of cyclin dependent kinase in chlamydomonas is required for coupling cell size to cell division. Elife 5:e10767. doi:10.7554/eLife.10767

Lyth O, Vizcay-Barrena G, Wright KE, Haase S, Mohring F, Najer A, Henshall IG, Ashdown GW, Bannister LH, Drew DR, Beeson JG, Fleck RA, Moon RW, Wilson DW, Baum J. 2018. Cellular dissection of malaria parasite invasion of human erythrocytes using viable Plasmodium knowlesi merozoites. Sci Rep 8:1–11. doi:10.1038/s41598-018-28457-z

Machado M, Steinke S, Ganter M. 2021. Plasmodium Reproduction, Cell Size, and Transcription: How to Cope With Increasing DNA Content? Front Cell Infect Microbiol 11:660679. doi:10.3389/fcimb.2021.660679

Mancio-Silva L, Lopez-Rubio JJ, Claes A, Scherf A. 2013. Sir2a regulates rDNA transcription and multiplication rate in the human malaria parasite Plasmodium falciparum. Nat Commun 4:1–6. doi:10.1038/ncomms2539

Mancio-Silva L, Slavic K, Grilo Ruivo MT, Grosso AR, Modrzynska KK, Vera IM, Sales-Dias J, Gomes AR, Macpherson CR, Crozet P, Adamo M, Baena-Gonzalez E, Tewari R, Llinás M, Billker O, Mota MM. 2017. Nutrient sensing modulates malaria parasite virulence. Nature 547:213–216. doi:10.1038/nature23009

Matthews H, Duffy CW, Merrick CJ. 2018. Checks and balances? DNA replication and the cell cycle in Plasmodium. Parasites and Vectors 11:216. doi:10.1186/s13071-018-2800-1

Mauritz JMA, Esposito A, Ginsburg H, Kaminski CF, Tiffert T, Lew VL. 2009. The Homeostasis of Plasmodium falciparum-Infected Red Blood Cells. PLoS Comput Biol 5:11–14. doi:10.1371/journal.pcbi.1000339

McDonald J, Merrick CJ. 2022. DNA replication dynamics during erythrocytic schizogony in the malaria parasites Plasmodium falciparum and Plasmodium knowlesi. PLoS Pathog 18:1–25. doi:10.1371/journal.ppat.1010595

McLean KJ, Jacobs-Lorena M. 2017. Plasmodium falciparum Maf1 confers survival upon amino acid starvation. MBio 8:e02317. doi:10.1128/mBio.02317-16

Mehnert AK, Simon CS, Guizetti J. 2019. Immunofluorescence staining protocol for STED nanoscopy of Plasmodium-infected red blood cells. Mol Biochem Parasitol 229:47–52. doi:10.1016/j.molbiopara.2019.02.007

Mohring F, Hart MN, Rawlinson TA, Henrici R, Charleston JA, Diez Benavente E, Patel A, Hall J, Almond N, Campino S, Clark TG, Sutherland CJ, Baker DA, Draper SJ, Moon RW, Benavente ED, Patel A, Hall J, Almond N, Campino S, Clark TG, Sutherland CJ, Baker DA, Draper SJ, Moon RW. 2019. Rapid and iterative genome editing in the malaria parasite Plasmodium knowlesi provides new tools for P. vivax research. Elife 1–29. doi:10.7554/elife.45829

Moon RW, Hall J, Rangkuti F, Ho Y, Almond N, Mitchell GH, Pain A, Holder AA, Blackman MJ. 2013. Adaptation of the genetically tractable malaria pathogen Plasmodium knowlesi to continuous culture in human erythrocytes. Proc Natl Acad Sci U S A 110:531–6. doi:10.1073/pnas.1216457110

Morahan BJ, Abrie C, Al-Hasani K, Batty MB, Corey V, Cowell AN, Niemand J, Winzeler EA, Birkholtz LM, Doerig C, Garcia-Bustos JF. 2020. Human Aurora kinase inhibitor Hesperadin reveals epistatic interaction between Plasmodium falciparum PfArk1 and PfNek1 kinases. Commun Biol 3:1–10. doi:10.1038/s42003-020-01424-z

Neumann FR, Nurse P. 2007. Nuclear size control in fission yeast. J Cell Biol 179:593–600. doi:10.1083/jcb.200708054

Ondracka A, Dudin O, Ruiz-Trillo I. 2018. Decoupling of Nuclear Division Cycles and Cell Size during the Coenocytic Growth of the Ichthyosporean Sphaeroforma arctica. Curr Biol 28:1964–1969.e2. doi:10.1016/j.cub.2018.04.074

Reilly Ayala HB, Wacker MA, Siwo G, Ferdig MT. 2010. Quantitative trait loci mapping reveals candidate pathways regulating cell cycle duration in Plasmodium falciparum. BMC Genomics 11:577. doi:10.1186/1471-2164-11-577

Reilly HB, Wang H, Steuter JA, Marx AM, Ferdig MT. 2007. Quantitative dissection of clone-specific growth rates in cultured malaria parasites. Int J Parasitol 37:1599–1607. doi:10.1016/j.ijpara.2007.05.003

Robbins JA, Absalon S, Streva VA, Dvorin JD. 2017. The Malaria Parasite Cyclin H Homolog PfCyc1 Is Required for Efficient Cytokinesis in Blood-Stage Plasmodium falciparum. MBio 8:e00605–17. doi:10.1128/mBio.00605-17

Rudlaff RM, Kraemer S, Marshman J, Dvorin JD. 2020. Three-dimensional ultrastructure of Plasmodium falciparum throughout cytokinesis. PLoS Pathog 16. doi:10.1371/journal.ppat.1008587

Simon CS, Funaya C, Bauer J, Voβ Y, Machado M, Penning A, Klaschka D, Cyrklaff M, Kim J, Ganter M, Guizetti J. 2021a. An extended DNA-free intranuclear compartment organizes centrosome microtubules in malaria parasites. Life Sci Alliance 4:e202101199. doi:10.26508/lsa.202101199

Simon CS, Stürmer VS, Guizetti J. 2021b. How Many Is Enough? - Challenges of Multinucleated Cell Division in Malaria Parasites. Front Cell Infect Microbiol 11:658616. doi:10.3389/fcimb.2021.658616

Waldecker M, Dasanna AK, Lansche C, Linke M, Srismith S, Cyrklaff M, Sanchez CP, Schwarz US, Lanzer M. 2017. Differential time-dependent volumetric and surface area changes and delayed induction of new permeation pathways in P. falciparum-infected hemoglobinopathic erythrocytes. Cell Microbiol 19:e12650. doi:10.1111/cmi.12650

White MW, Suvorova ES. 2018. Apicomplexa Cell Cycles: Something Old, Borrowed, Lost, and New. Trends Parasitol 34:759–71. doi:10.1016/j.pt.2018.07.006

WHO. 2021. WHO Global, World Malaria Report 2021, Word Malaria report Geneva: World Health Organization. (2021). Licence: CC.

Yahiya S, Jordan S, Smith HX, Gaboriau DCA, Famodimu MT, Dahalan FA, Churchyard A, Ashdown GW, Baum J. 2022. Live-cell fluorescence imaging of microgametogenesis in the human malaria parasite Plasmodium falciparum. PLoS Pathog 18. doi:10.1371/journal.ppat.1010276

Zigler JS, Lepe-zuniga B Vistica JL, Gery I. 1985. Analysis of the cytotoxic effects of light-exposed hepes-containing culture medium. Vitr Cell Dev Biol 21:282.

