## Supplemental information for "Progeny counter mechanism in malaria parasites is linked to extracellular resources"

**Contents**

Figures S1 to S9

Movie captions S1 to S5

Supplemental movies can be accessed under: https://www.dropbox.com/sh/dsnbp76ofq7see7/AAD66NmUsU2bA04CE7I7vKfqa?dl=0

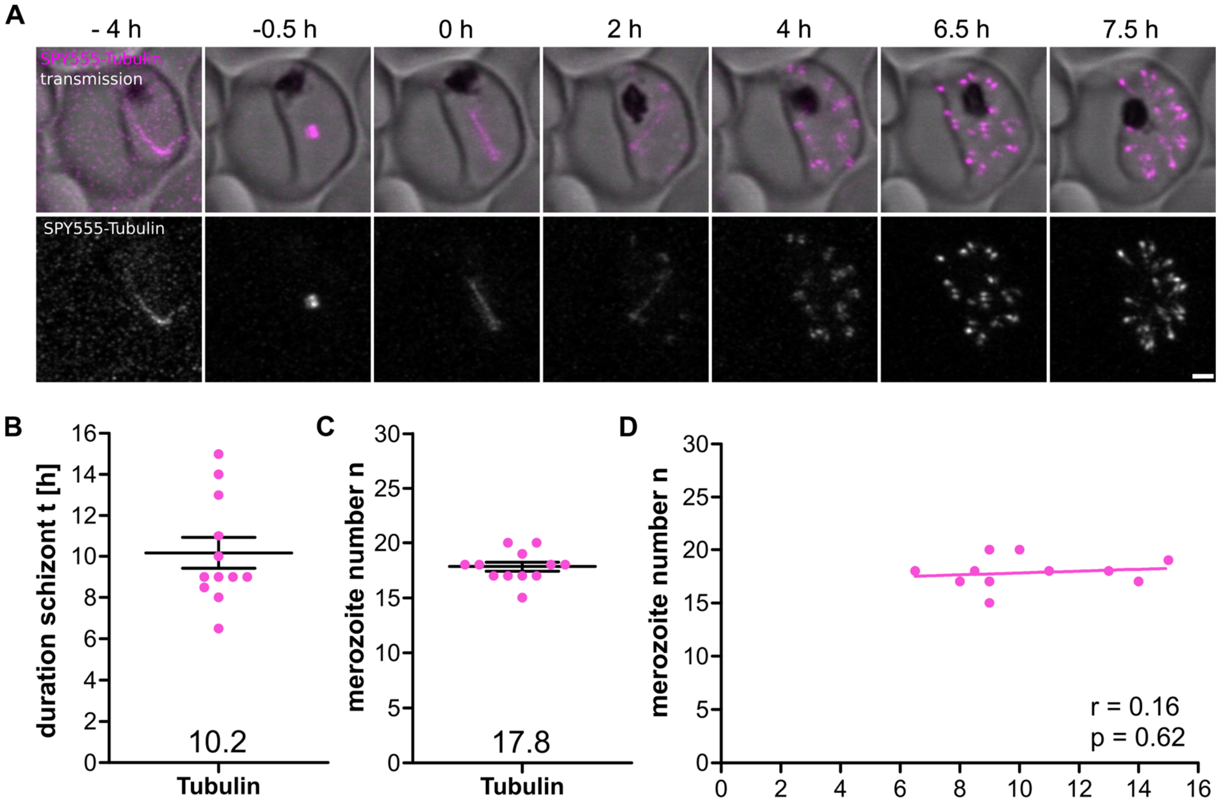


***Supplemental Figure 1. Quantification of schizogony duration and number of daughter cells using microtubule maker in P. falciparum is reproducible. A*** *Airyscan-processed time-lapse images of P. falciparum strain 3D7 stained with SPY555-Tubulin. Timepoint of first spindle elongation was set to 0 h and the timepoint where segmentation started was considered as schizogony end. Shown are maximum intensity projections. Scale bar is 1 µm.* ***B*** *Quantification of schizont stage duration in hours* ***C*** *final merozoite number based on number of subpellicular microtubule structures. Black bar represents mean and SEM.* ***D*** *Correlation of merozoite number against duration of schizont stage. Given are Pearson correlation coefficient r and p values. N = 12.*

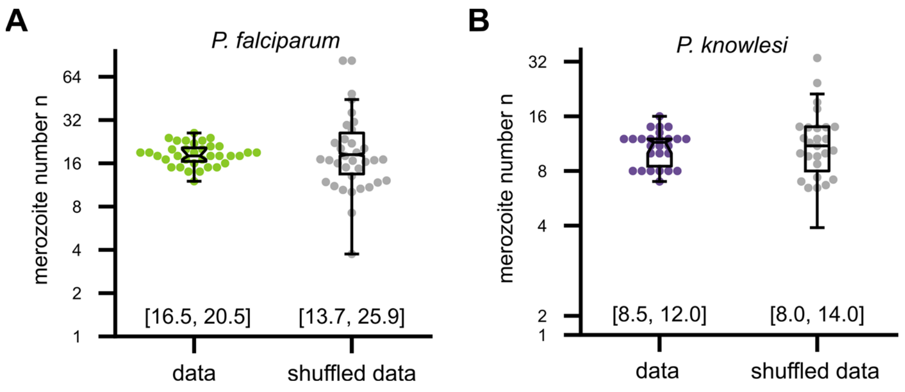


***Supplemental Figure 2. Variability in merozoite number is reduced by the negative correlation between duration of schizont stage t and multiplication rate λ. A*** *Merozoite number data from Fig. 1D alongside reshuffled data generated by breaking up t and λ pairs. Shown are SD and quartiles. Shuffled data points represent a swarm plot. Variation of shuffled data is higher as indicated by the interquartile ranges below.* ***B*** *as in A for the* P. knowlesi *data.*

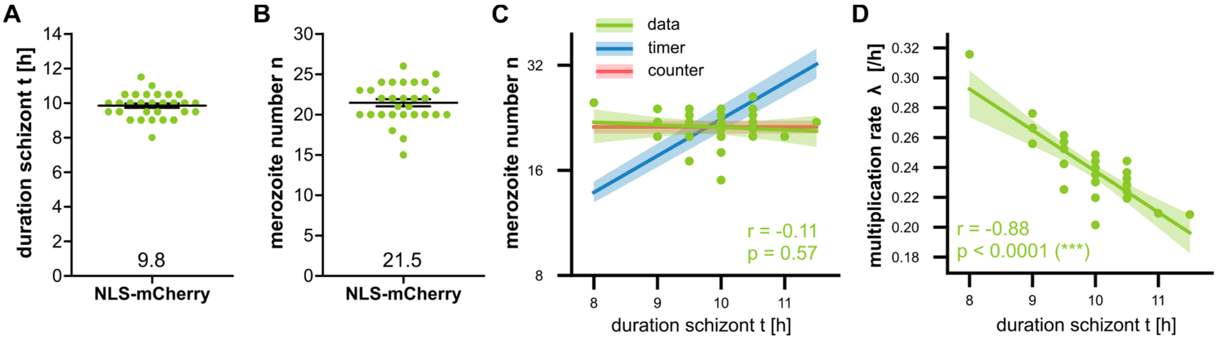


***Supplemental Figure 3. Quantification of duration of schizont stage and merozoite number in P. falciparum expressing nuclear mCherry yields values consistent with H2B-GFP strain. A** Quantification of schizont stage duration in hours and **B** merozoite number of P. falciparum 3D7 ectopically expressing a nuclear mCherry signal (NLS-mCherry). Given are means and SEM. **C** Plot comparing predicted regression curves for timer (blue) and counter (red) model with data bootstrapped with 95% confidence interval. **D** Bootstrapped regression curves of t against λ. N = 29 all from three independent replicas. Given are Pearson correlation coefficient r and p values.*

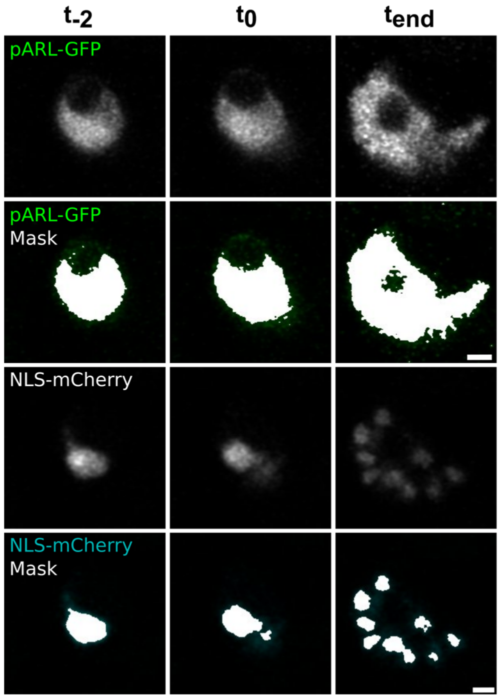


***Supplemental Figure 4. Image segmentation of cytosolic and nucleoplasmic markers allows for volume calculations.*** *Exemplary single image slices from time points pre-schizogony ($t_{-2}$), schizogony start ($t_0$) and schizogony end ($t_{end}$) showing cytoplasmic marker pARL-GFP (green), nuclear marker NLS-mCherry (cyan), and thresholding masks generated for measuring respective areas and volumes. GFP mask was determined by automatic thresholding and mCherry by manual and visual adjustment. Shown is z slice 11 of a stack of 20 slices. Contrast was adjusted for better visualization. Scale bar is 1 µm.*

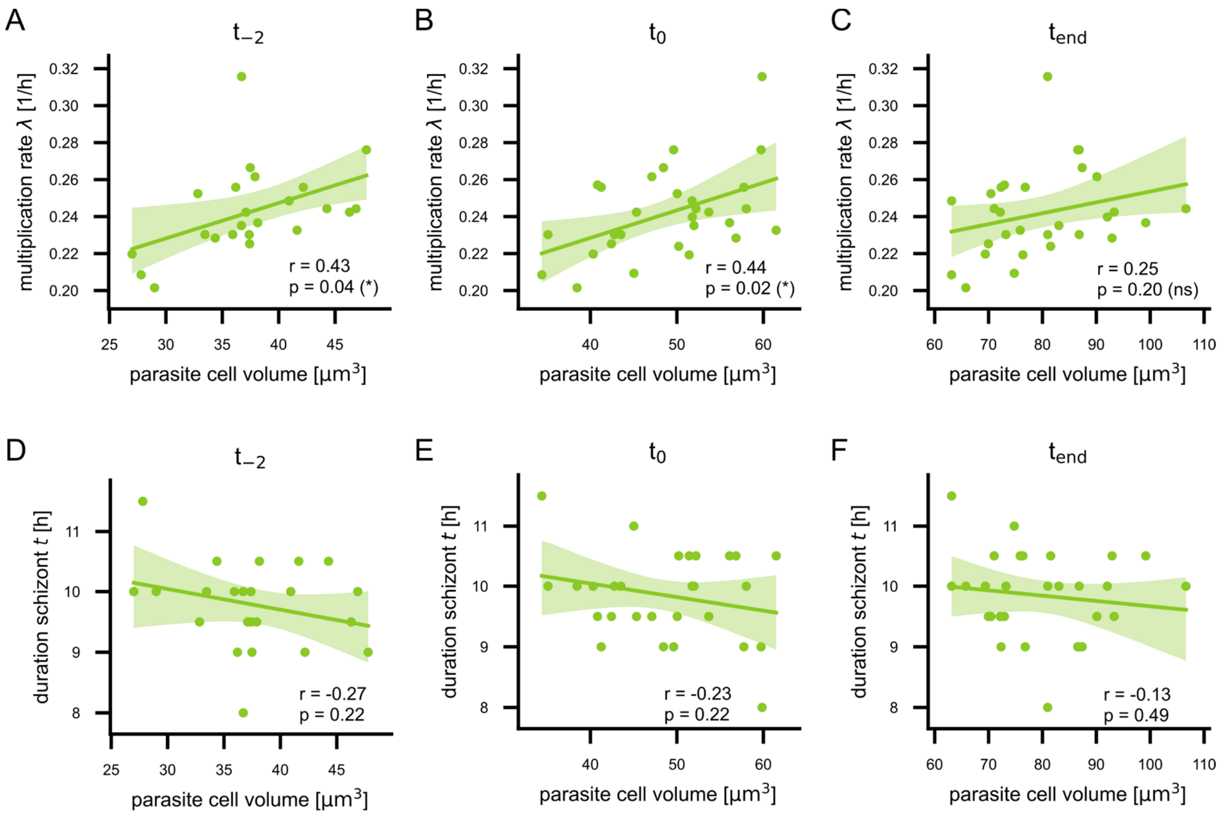


***Supplemental Figure 5. Correlations of t and λ against parasite cell volume at three timepoints of schizogony.*** *Show are regression curves of cellular parameters t and λ measured in P. falciparum 3D7 ectopically expressing a nuclear mCherry signal and cytoplasmic GFP plotted against parasite cell volume measured at replication start, schizont stage onset, and schizont stage end. N = 23 for t-2 and N = 29 for all others from three independent replicates. Given are Pearson correlation coefficient r and p values. Values are bootstrapped to 95% confidence interval.*

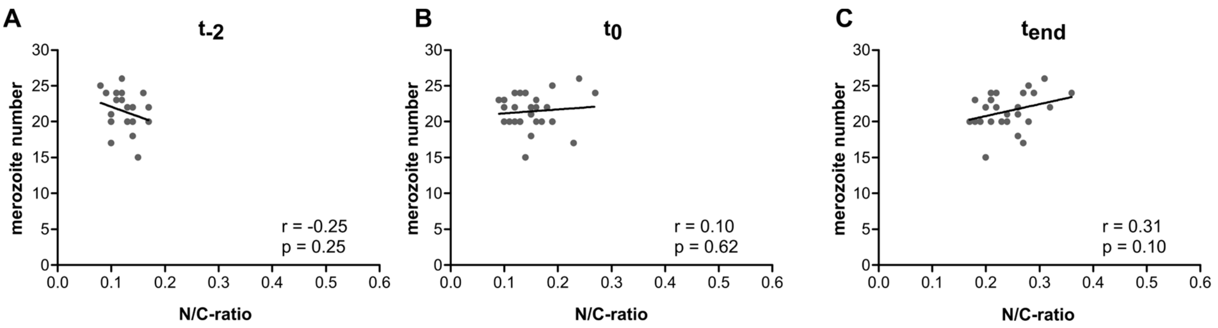


***Supplemental Figure 6. Merozoite number does not correlate with N/C-ratio**. Correlation of merozoite number against N/C ratio all for **A** pre-schizogony ($t_{-2}$), **B** schizogony start ($t_0$) and **C** schizogony end ($t_{end}$). Given are Pearson correlation coefficient r and p values. N = 23 for $t_{-2}$ and N = 29 for all others.*

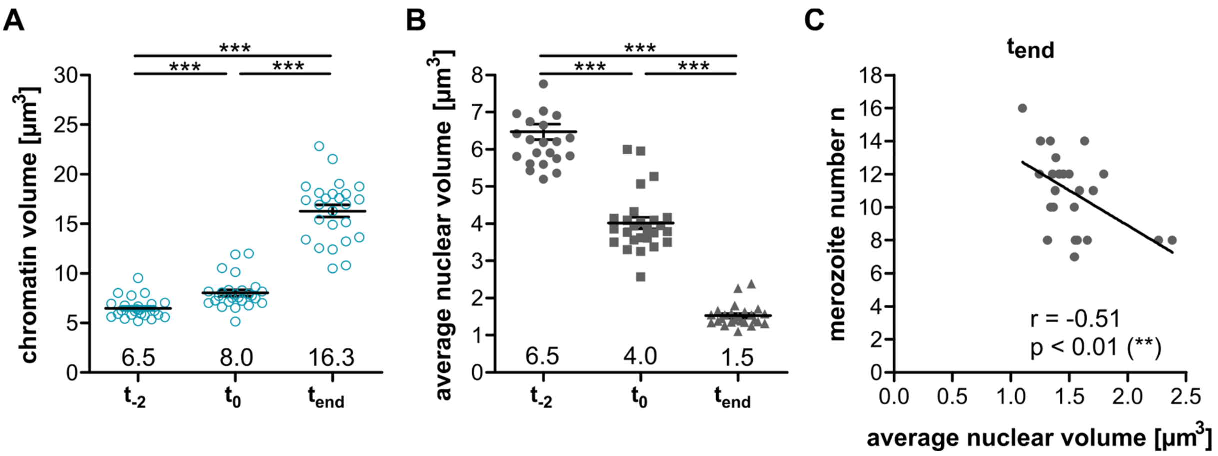


***Supplemental Figure 7. P. knowlesi compacts its chromatin during schizogony. A*** *Chromatin volumes based on H2B-GFP marker for individual cells at three timepoints; pre-schizogony ($t_{-2}$), schizogony start ($t_0$) and schizogony end ($t_{end}$). Error bars represent mean and SEM. Statistics: t-test with Welch's correction.* ***B*** *Average nuclear volume (total chromatin volume divided by nuclear number) for individual cells.* ***C*** *Correlation of merozoite number against average nuclear volume at end of schizogony ($t_{end}$). Given are Pearson correlation coefficient r and p values. N = 24.*

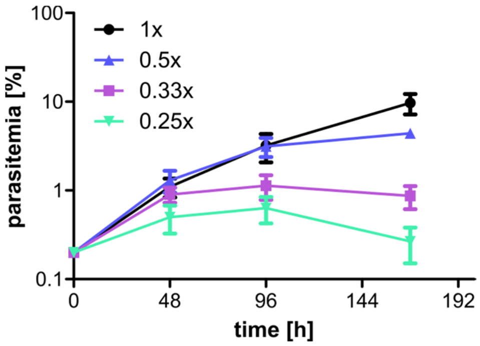


***Supplementary Figure 8. Parasites can be cultured in medium diluted to 0.5x.*** *Growth curve with asynchronous parasite cultures cultivated in normal (1x, black) and diluted (0.5x, blue; 0.33x, purple; 0.25x, green) medium with 0.9% NaCl. Medium was changed every 48h and parasitemia was assessed using Giemsa-stained thin blood smears. Plotted are mean and SD of three technical replicates.*

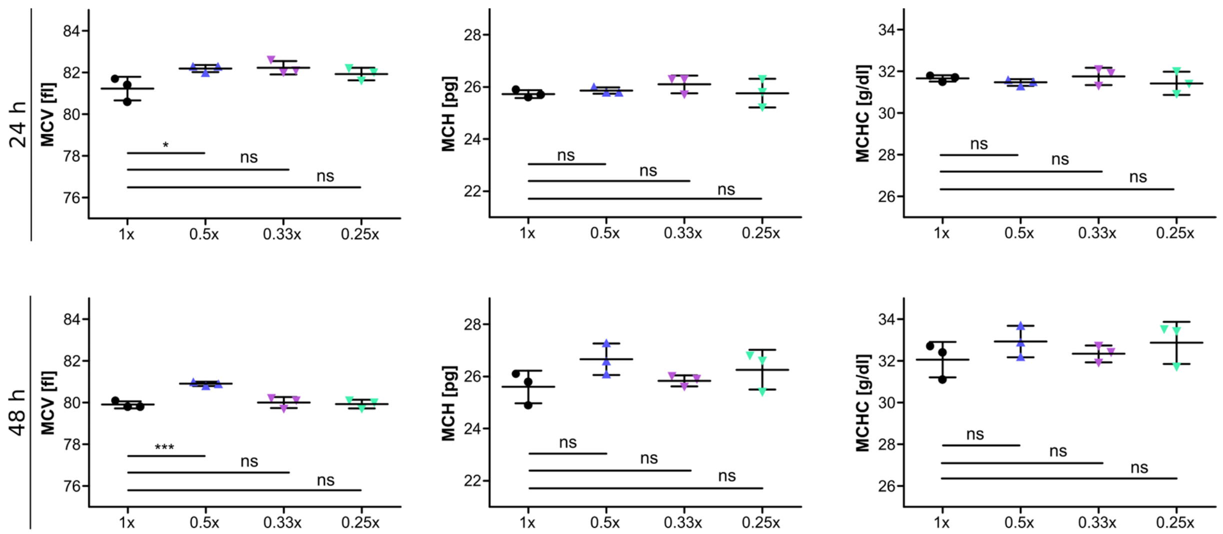


***Supplementary Figure 9. Medium dilution has no major effect on red blood cell indices.*** *Automated haematology analyses of erythrocytes after 24 and 48 h in different medium dilutions (1x, black; 0.5x, blue; 0.33x, purple; 0.25x, green). Plotted are the indices mean cell volume (MCV) in fl, mean cell hemoglobin (MCH) in pg and mean cell hemoglobin concentration (MCHC) in g/dl. Shown are mean and SD of three technical replicates.*

***Movie S1 H2B-GFP P. falciparum.*** *Airyscan-processed time-lapse imaging of P. falciparum 3D7 ectopically expressing PfH2B tagged with GFP. Shown are images taken every hour during a total of 14 hours as maximum intensity projections of 21 slices in a 0.3 µm interval. Left, GFP; right, Brightfield and GFP (green). Image size is 15 x 15 µm.*

***Movie S2 H2B-GFP P. knowlesi.*** *Airyscan-processed time-lapse imaging of P. knowlesi A1-H.1 ectopically expressing PkH2B tagged with GFP. Shown are images taken every 30 min during a total of 13.5 hours as maximum intensity projections of 20 slices in a 0.36 µm interval. Left, GFP; right, Brightfield and GFP (green). Image size is 12 x 12 µm.*

***Movie S3 SPY555-Tubulin P. falciparum****. Airyscan-processed time-lapse imaging of P. falciparum 3D7 stained with live Tubulin dye (SPY555-Tubulin, 1:2000). Shown are images taken every 30 min during a total of 14.5 hours as maximum intensity projections of 13 slices in a 0.5 µm interval. Left, Tubulin; right, Brightfield and Tubulin (magenta). Image size is 12 x 12 µm.*

***Movie S4 pARL-GFP NLS-mCherry P. falciparum.*** *Airyscan-processed time-lapse imaging of P. falciparum 3D7 ectopically expressing cytosolic GFP (pARL-GFP) and nuclear mCherry (NLS-mCherry). Shown are images taken every 30 min during a total of 17 hours as maximum intensity projections of 20 slices in a 0.36 µm interval. Left, pARL-GFP (green), NLS-mCherry (magenta); right, Brightfield, pARL-GFP, and NLS-mCherry. Image size is 12 x 12 µm.*

***Movie S5 pARL-GFP NLS-mCherry P. falciparum in 0.5x diluted culture medium.*** *Airyscan-processed time-lapse imaging of P. falciparum 3D7 ectopically expressing cytosolic GFP (pARL-GFP) and nuclear mCherry (NLS-mCherry). Acquisition started after about 24 hours incubation in diluted medium. Shown are images taken every 30 min during a total of 17 hours as maximum intensity projections of 20 slices in a 0.36 µm interval. Left, pARL-GFP (green), NLS-mCherry (magenta); right, Brightfield, pARL-GFP, and NLS-mCherry. Image size is 11 x 11 µm.*
